# Concomitant Pyroptotic and Apoptotic Cell Death Triggered in Monocytes Infected by Zika Virus

**DOI:** 10.1101/2021.09.02.458664

**Authors:** Chunxia Wen, Yufeng Yu, Chengfeng Gao, Xian Qi, Carol J. Cardona, Zheng Xing

## Abstract

Zika virus (ZIKV) is a positive-sense RNA flavivirus and can cause serious neurological disorders including microcephaly in infected fetus. As a mosquito-borne arbovirus, ZIKV enters bloodstream and is transmitted into the fetus through the placenta in pregnant women. Monocytes are considered one of the earliest blood cell types to be infected by ZIKV. As a first line defence, monocytes are crucial components in innate immunity and host responses and may impact viral pathogenesis in humans. Previous studies have shown that ZIKV infection can activate inflammasomes and induce proinflammatory cytokines in monocytes. In this report, we showed that ZIKV carried out a productive infection, which lead to cell death in human and murine monocytic cells. In addition to the presence of cleaved caspase-3, indicating that apoptosis was involved, we identified the cleaved caspase-1 and gasdemin D (GSDMD) as well as increased secretion of IL-1β and IL-18, suggesting that the inflammasome was activated that may lead to pyroptosis in infected monocytes. The pyroptosis was NLRP3-dependent and could be suppressed in the monocytes treated with shRNA to target and knockdown caspase-1, or an inhibited for caspase-1, indicating that the pyroptosis was triggered via a canonical approach. Our findings in this study demonstrate a concomitant occurrence of apoptosis and pyroptosis in ZIKV-infected monocytes, with multiple mechanisms involved in the cell death, which may have potentially significant impacts on viral pathogenesis in humans.

## INTRODUCTION

Zika virus (ZIKV) is a member of *Flaviviridae* family, which includes a large group of viruses that cause West Nile encephalitis, Dengue Fever, Japanese encephalitis, Tick-borne encephalitis and other important human diseases (1). ZIKV infection is usually self-limited in most of cases, which are either asymptomatic or mild with only fever, rash, conjunctivitis and malaise. ZIKV has been associated with Guillain-Barre syndrome or other mild neurological symptoms in some adults (2) and caught the world attention when it was linked to congenital infections, leading to spontaneous abortions and severe neonatal birth defects including microcephaly when a severe outbreak occurred in South America in late 2015 to 2016 (3).

Innate immunity plays a critical role in the early phase of viral infections and host defense, in which monocytes and macrophages, originated from bone marrow myeloid progenitor cells are key players (4). Once an infection occurs, monocytes activate their phagocytic function and release a variety of cytokines and chemokines, which will further promote their activation and differentiation (5, 6). Monocytes can become macrophages when they egress from the bloodstream and reside into tissues and organs via chemotaxis. In addition to cytokine and chemokine release, monocytes/macrophages recruit lymphocytes and activate adaptive immunity through antigen presentation (7) and help clear viral infection in the host. On the other hand, monocytes infected with viruses are considered possible to serve as a Trojan horse under certain circumstances that promotes virus spread and leads to virus dissemination within the host. More importantly, this could happen to bring viruses into the immune privileged tissues and organs such as the placenta, testes, and brain when monocytes migrate across protective blood barriers in the host (8). Indeed several studies indicated that monocytes could severe as virus reservoirs and facilitate virus dissemination and transmigration into the brain through blood-brain barrier (BBB) (9, 10).

Despite the viremia in blood of Zika patients, knowledge about the exact target cells and their responses in the blood during ZIKV infection remains limited. Blood CD14+ monocytes appears to be the primary cells for ZIKV infection, which leads to differential immunomodulatory response or M2-skewed immunosuppression during pregnancy (11). A recent report shows that monocytes exhibit more adhesion molecules and abilities to attach onto the vessel wall and transmigrate across endothelia which promotes ZIKV dissemination to neural cells (12). Several studies have shown that ZIKV infection can activate the NLRP3 inflammasome, which results in the secretion of pro-inflammatory cytokines (13-15). These findings make it complicated to assess the role of monocytes in ZIKV infection, and in particular in viral pathogenesis in an infection during pregnancy. However, no study has shown what the fate is for the infected monocytes or whether these cells die of pyroptosis due to the inflammasome activation (13-16), while placental macrophages appear to be resistant to cell death during ZIKV infection (17).

Host cells can react by activating various innate defenses in response to viral infections. In addition to antiviral or pro-inflammatory cytokines and chemokines, cells could trigger programed cell death with complex outcomes, which may eliminate infected cells and clear virus replicative niche (18). Apopotosis and pyroptosis are caspase-dependent cell death and necroptosis is activated relying on the activation of phosphorylated receptor interacting serine/threonine-protein kinase (RIPK)(19-23). In this report, we showed the evidence that a productive ZIKV infection led to cell death in both human and murine monocytes. Our data indicated that the infected monocytes died of apoptosis and pyroptosis with the presence of cleaved caspase-1 and caspase-3 and processed GSDMD. The pyroptosis in human and murine monocytes was dependent on the NLRP3 inflammasome activation induced by ZIKV infection. Although previous studies have indicated the significance of the inflammasomes in proinflammatory cytokine responses(13-16) and inhibition of the cGAS-mediated interferon signaling in ZIKV monocytes (13), concomitant apoptosis and pyroptosis may benefit the host by removing a virus shelter and preventing viral spreading and disseminating, which could be of significance to viral pathogenesis in human ZIKV infection.

## MATERIALS AND METHODS

### Cell Lines and Cultures

THP-1 cells and human embryonic kidney cells (HEK293T) were purchased from the Cell Bank of Chinese Academy Sciences (Shanghai, China). RAW264.7 cells were purchased from the American Type Culture Collection (ATCC). RAW264.7 and HEK293T cells were cultured in Dulbecco’s modified Eagle’s medium (DMEM) (Gibco, Grand Island, NY) supplemented with 10% fetal bovine serum (FBS, ExCell Bio., China), 100 U/ml penicillin, and 100 μg/ml streptomycin sulphate (Beyotime Biotechnology, China). Cells were cultured in an incubator at 37°C with a humidified atmosphere of 5% CO_2_.

### Reagents and Antibodies

ZIKV-E antibody (#B1845) was purchased from Beijing Biodragon Immunotech (Beijing, China). Antibodies for pro-caspase-1 antibody (#ab179515), phospho-MLKL (#ab187091), phospho-RIPK3 (#209384) and GSDMD (#ab210070) were purchased from Abcam (Cambridge, MA). Another GSDMD antibody (#A10164) was purchased from ABclonal. Antibodies for NLRP3 (#D4D8T), pro-caspase-3 (#9555S), cleaved caspase-3 (#9664S), pro-PARP (#9532S), cleaved-PARP (#9541S), and phospho-RIPK1 (#44590S) were purchased from Cell Signalling Technology (Beverly, MA). We also purchased antibodies for cleaved caspase-1 (#AF4022) from Affinity Biosciences (Taizhou, China) and GAPDH and β-actin from Proteintech (Wuhan, China). An MTT Assay Kit was purchased from SunShineBio (Nanjing, China). ELISA kits for mouse interleukin-1β, mouse interleukin-18, human interleukin-1β, and human interleukin-18 were purchased from BOSTER (Wuhan, China). Compounds VX765, ZVAD-FMK, GSK’872, Nec-1s, and Phorbol-12-myristate-13-acetate (PMA) were purchased from Selleck. RNAiso plus reagent was purchased from TAKARA.

### Virus Infection

In this study ZIKV SZ01 strain (GenBank: KU866423), a gift from Dr Shibo Jiang of Fudan University, was used. THP-1 cells were activated with 100 nM PMA for 24 to 48hrs (24). THP-1 and RAW264.7 cells were inoculated with 1 MOI of ZIKV SZ01 for infection at 37°C for 12 to 48 hrs. Cell medium was collected and cell lysates or total RNA were prepared for further analyses.

### MTT Assay for Cell Viability

PMA-activated THP-1 cells and RAW264.7 cells were inoculated with ZIKV (0.01, 0.1 or 1 MOI) for 12, 24, 36 and 48 hrs prior to addition of 3-(4,5)-dimethylthiahiazo (-z-y1)-3,5-di phenytetrazoliumromide (MTT) for another 4 hrs. Cell viability was determined as a ratio of absorbance at OD_570_ of ZIKV-infected cells to uninfected cells. The assay was carried out at least three times for each group. Unpaired Student’s t-test was used to evaluate the data. The data shown are the mean ± SD of three independent experiments. *P<0.05, ***P< 0.001.

### Quantitative Realtime PCR

Total RNA was extracted with RNAiso plus reagent for quantitative realtime PCR (QPCR) following the manufacture’s manual. QPCR was performed with 1μl of cDNA in a total volume of 20μl with SYBR Green QPCR Master Mix (Vazyme) according to the manufacturer’s instructions. QPCR primers were designed by Primer Premier 5.0. Relative gene expression levels were normalized by β-actin housekeeping gene. Relevant fold change of each gene was calculated by following the formula: 2^ΔCt of gene-ΔCt of β-actin^ (ZIKV infected cells)/ 2^ΔCt of gene-ΔCt of β-actin^ (ZIKV - uninfected cells).

### Lentivirus Packaging for shRNA

Lentivirus vectors were obtained from Shanghai Jiao Tong University. The negative control was a pLKO.1 vector containing sequences encoding shRNA. To ensure knockdown efficiency, we selected three shRNA sequences for each targeted gene. The sense sequences for pro-caspase-1 shRNA are: CTCTCATTATCTGCAATGA, AGCGTAGATGTGAAAAAAA, and CCAGATATACTACAACTCA. The sense sequences for NLRP3 shRNA are: TCGAGAATCTCTATTTGTA, ACGCTAATGATCGACTTCA, and AGGAGAGACCTTTATGAGA. PMA-activated THP-1 cells were infected with the recombinant lentiviruses expressing shRNAs targeting pro-caspase-1 or NLRP3. 48 hrs later, the culture medium was discarded and the cells were inoculated with ZIKV at 1 MOI. The culture medium and cell lysates were harvested at indicated time points for enzyme-linked immunosorbent assay (ELISA) and western blot analyses.

### ELISA and Western Blot Analysis

Secretion of cytokine IL-1β and IL-18 in culture medium was measured using commercial ELISA kits. Each group of testing was replicated for three times and resultant data were analysed by Student’s t-test. Cell lysates from the THP-1 (wild type, knockdown cells for NLRP3 and pro-caspase-1) and RAW264.7 (wild type and knockout cells for NLRP3) cells were prepared by RIPA lysis buffer with 1% PMSF. Protein concentrations were determined by a Bradford assay (BCA). Cell lysates (40 μg) were electrophoresed in 10-15% SDS-PAGE and the proteins transferred to a PVDF membrane for subsequent western blot analyses. Protein signals on the membrane were visualized by a GelCap ECL analyzer (Canon).

### Confocal Immunofluorescence

ZIKV-infected THP-1 or RAW264.7 cells on chamber slides were fixed by 4% paraformaldehyde at room temperature for 15 min and permeabilized with 0.1% Triton-X-100 for 10 min. The cells were washed three times with PBS and then blocked with 5% BSA for 1 hr at room temperature. The cells were then incubated with an antibody for GSDMD overnight at 4°C, followed by four washes with PBS. The slides were incubated with Alexa Fluor 594-conjugated Affinipure goat-anti-rabbit IgG (H+L) (Proteintech) for 1hr at room temperature. After four more washes, the cells were incubated with DAPI solution for 10 min to stain the nuclei. The cells were finally analysed under a confocal laser scanning microscope (FV3000, Olympus).

### Statistical Analysis

Student’s t-test was used to evaluate the data. The data shown are the mean ±SD of three independent experiments. The differences with a value of p<0.05 were considered statistically significant.

## RESULTS

### Infection of ZIKV in Monocytes Triggered Cell Death

As an arbovirus, ZIKV enters bloodstream and may infect and replicate in some blood cells. We chose monocytes, which are susceptible to ZIKV as reported previously, for infection to examine the ultimate fate of the infected cells. Two monocytic cell lines, THP-1 (human) and RAW264.7 (mouse), were pre-treated with PMA, followed by inoculation of ZIKV virus at 1 MOI. Morphological changes were observed under a microscope, which showed that the cells became swelling, detached, and their membrane deteriorated. Eventually the cell body broke up into debris after 24 or 48 hrs post infection (p.i.) (Figure 1A-B). The cell death caused by ZIKV infection was dose dependent. When the cells were infected with the virus at MOIs of 0.01, 0.1, or 1, cell death occurred at various time points p.i. and more death was observed in the culture infected with higher amount of viral doses (Figure 1C-D). To confirm that a productive infection occurred, we detected viral RNA replication which showed that viral E gene copy numbers increased over time p.i. by realtime RT-PCR in both THP-1 and RAW264.7 cells (Figure 1E-F). Viral E protein could also be detected starting mainly at 12 hrs p.i. in the infected cells (Figure 1G-H).

**Figure 1.**
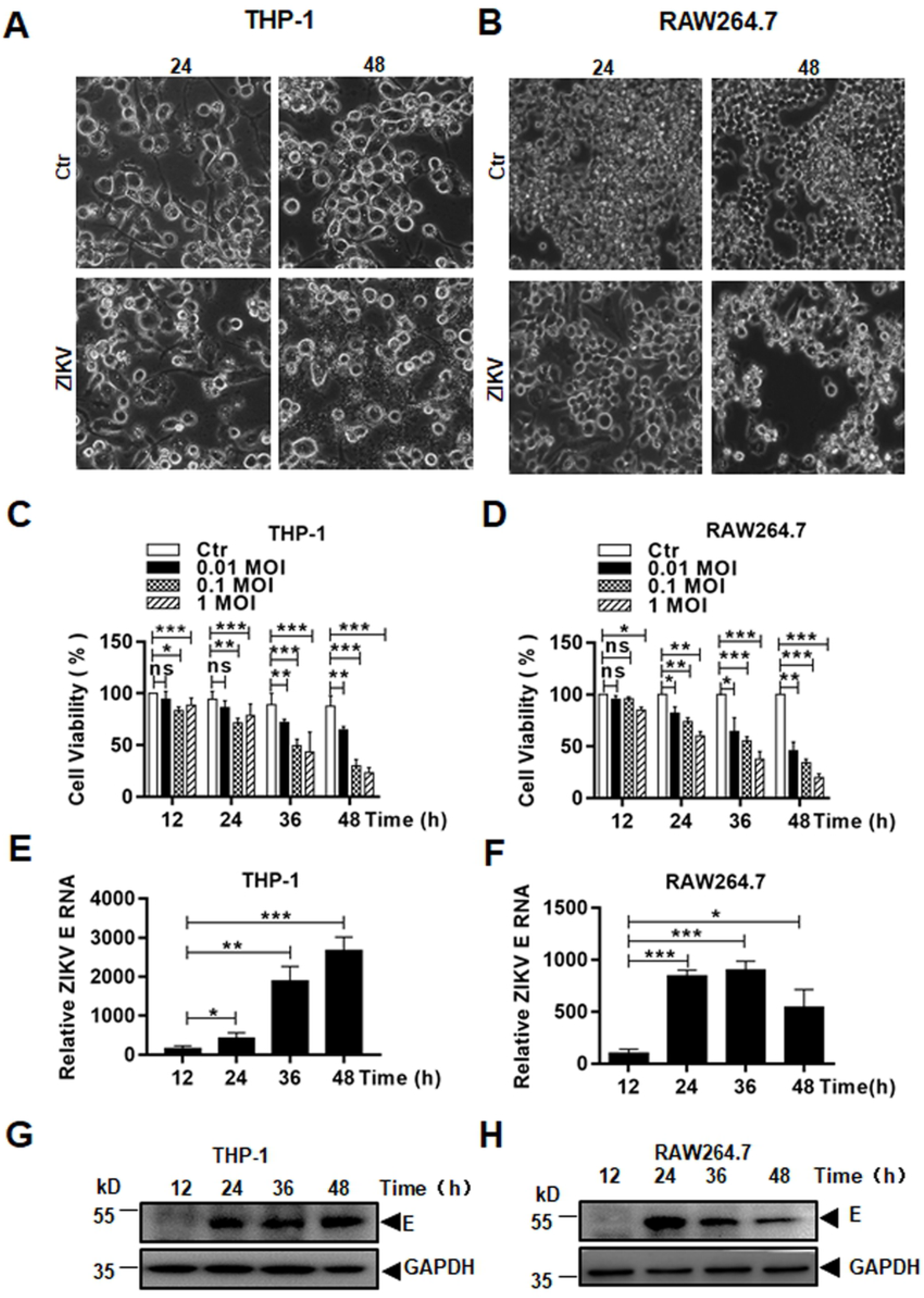
Viral replication and cell death caused in ZIKV-infected monocytic cells. Human and mouse monocytic THP-1 (**A**) and RAW264.7 (**B**) cells were infected with ZIKV at 1 MOI and shown cytopathic effect (CPE) at 24 and 48 hrs p.i. (Magnification x 40). Cell viability was assessed at various time points p.i. in THP-1 (**C**) and RAW264.7 (**D**) cells infected with ZIKV at various MOIs by an MTT assay. Total RNA was prepared for realtime RT-PCR to quantify copy numbers of ZIKV E RNA in infected THP-1 (**E**) and RAW264.7 (**F**) ZIKV E protein expression was detected in cell lysates prepared from infected THP-1 (**G**) and RAW264.7 (**H**) cells by western blot analyses. The data were presented as mean ±SD and analysed by Student’s t-test. *, P<0.05; **, P<0.01; ***, P< 0.001.

### Programed Cell Death Induced in Monocytes with ZIKV Infection

We tried to understand the mechanism about how the monocytes died in response to ZIKV infection. THP-1 and RAW264.7 cells were pre-treated with ZVAD-FMK (40 μM), Nec-1s (50 μM), and GSK’872 (20 nM), the inhibitors for pan-caspases, RIPK1, and RIPK3, respectively, followed by infection with ZIKV. First, we observed the cell morphological changes in infected cells with or without the treatment of inhibitors. In the cells untreated with the inhibitors, the infected cells underwent cell death. It appeared that ZVAD-FMK could relieve the cell death in both THP-1 and RAW264.7 cells, although Nec-1s or GSK’872 had no apparent impact on the cell death at 24 and 48 hrs p.i. (Figure 2A-B), indicating that caspases could be involved in the cell death. We quantified the cell death by measuring cell viabilities with an MTT assay. The treatment with ZVAD-FMK significantly relieved the monocytes from cell death at 24 and 48 hrs p.i. in both THP-1 (Figure 3A) and RAW264.7 (Figure 3B) cells. On the other hand, the treatment with Nec-1s did not relieve the cells from death until 48 hrs p.i. in THP-1 cells and had no effect on cell death in RAW264.7 cells; the treatment with GSK’872 did not reverse cell death in both THP-1 and RAW264.7 cells. Taken together, these data suggest that caspases, but not RIPKs, may play a role in the cell death triggered by ZIKV infection.

**Figure 2.**
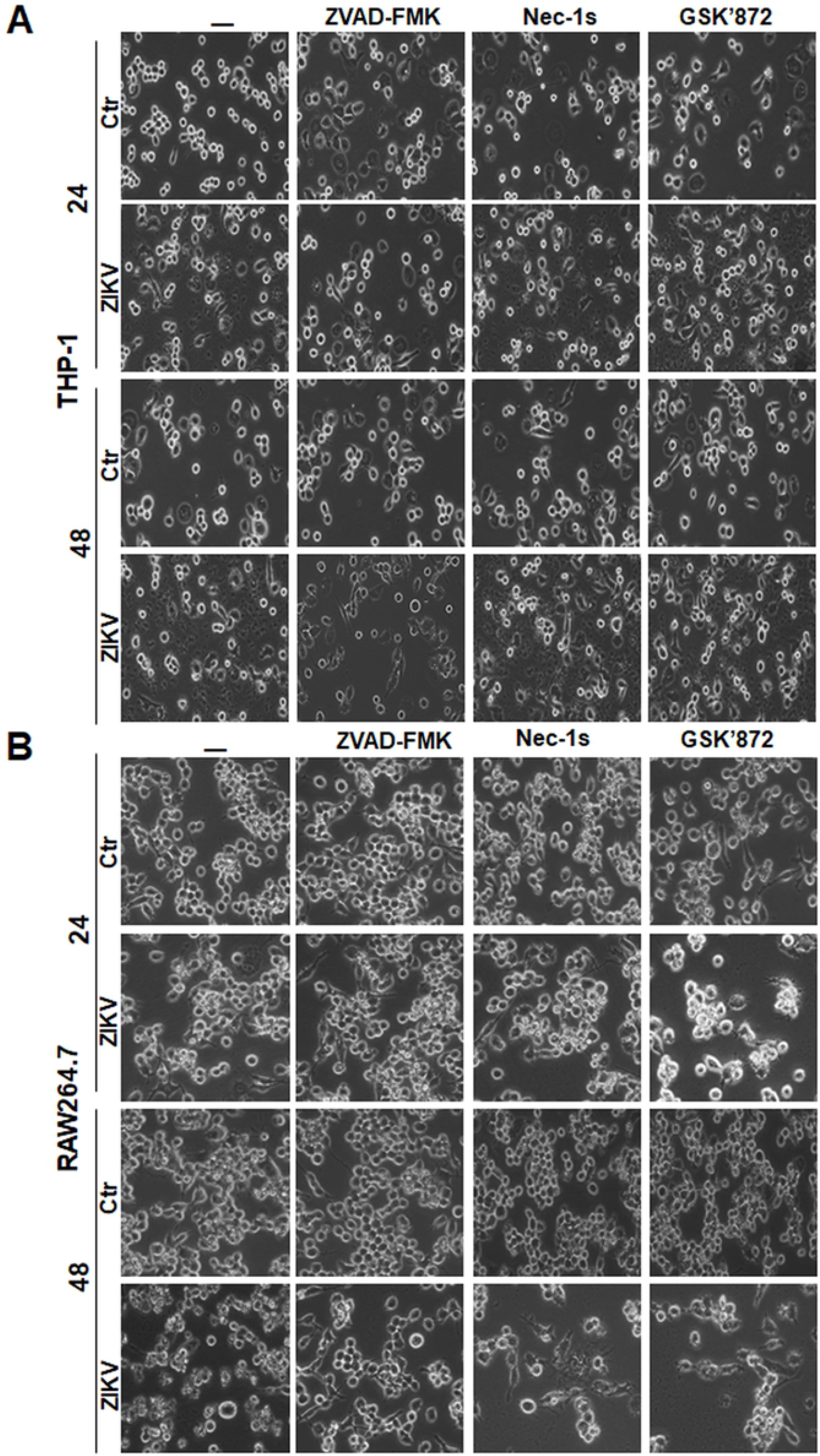
Morphological changes of cell death induced in ZIKV-infected monocytes pretreated by programmed death inhibitors. THP-1 (**A**) and RAW264.7 (**B**) cells were pretreated with pan-caspases inhibitor, ZVAD-FMK, or RIPK inhibitors, Nec-1s and GSK’872, followed by infection with ZIKV at 1 MOI. The cells were observed at 24 and 48 hrs p.i. under a light microscope (Magnification x 40).

**Figure 3.**
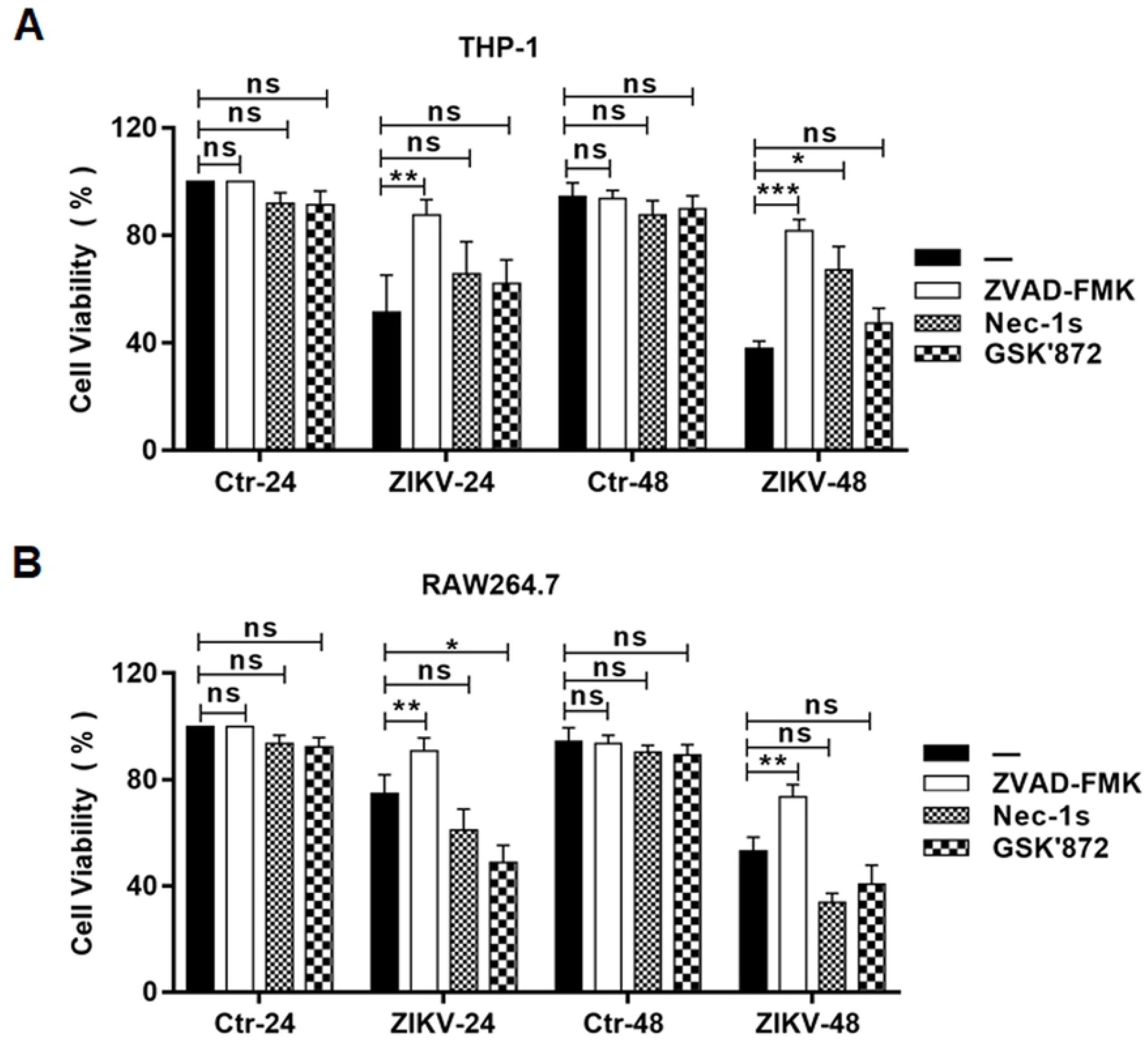
Blockage of cell death induced in ZIKV-infected cells by the inhibitors of programmed cell death. THP-1 (**A**) and RAW264.7 (**B**) cells were pre-treated with pan-caspases inhibitor, ZVAD-FMK, or RIPK inhibitors, Nec-1s and GSK’872, followed by infection with ZIKV at 1 MOI. The cell viability was assessed at 24 and 48 hrs p.i. with an MTT assay. The data were shown as mean ±SD and analysed by unpaired Students t-test. *, P<0.05; **, P<0.01; ***, P<0.001; ns, no significance.

We analysed the cell lysates prepared at various time points p.i. from the infected cells for western blot analyses. As shown in supplemental Figure S1, pro-caspase 3 was cleaved and activated, together with the presence of the cleaved substrate PARP, at the early stage of infection, confirming that apoptosis was activated in ZIKV-infected monocytes. We could also detect increased phosphorylation of RIPK1 and RIPK3, as well as phosphorylated MLKL, which indicates an activation of necroptosis. However, the activation of RIPKs may not have efficiently triggered a programed necrosis. These data suggest that caspases-dependent programed cell death could be the main cause involved in ZIKV-infected monocytes.

### Pyroptosis was induced in both human and murine monocytes during ZIKV infection

Considering that caspases are involved in not only apoptosis but also pyroptosis, we decided to examine what types of caspases were required for the cell death in ZIKV-infected monocytes. Pro-caspase-1 was activated as shown in the early studies indicating that inflammasomes were activated in monocytes. We confirmed that in both THP-1 and RAW264.7 cells (Figure 4A-B) pro-caspase-1 was processed and cleaved caspase-1 was detected at various time points p.i. by western blot analyses. In fact, expression of pro-caspase-1 was upregulated in both cell lines after ZIKV infection. In addition, GSDMD, a substrate of caspase-1, was cleaved to become cleaved GSDMD, an executor of pyroptosis, indicating that ZIKV infection activated inflammasomes, and induced pyrotosis in monocytes, probably via a caspase-1-dependent canonical pathway. Quantitative analyses showed the significant pro-caspase-1 upregulation (Figure 4C-D) and increases of cleaved caspase-1 (Figure 4E-F) and GSDMD (Figure 4 G-H) in infected THP-1 and RAW264.7 cells than in the cells uninfected.

**Figure 4.**
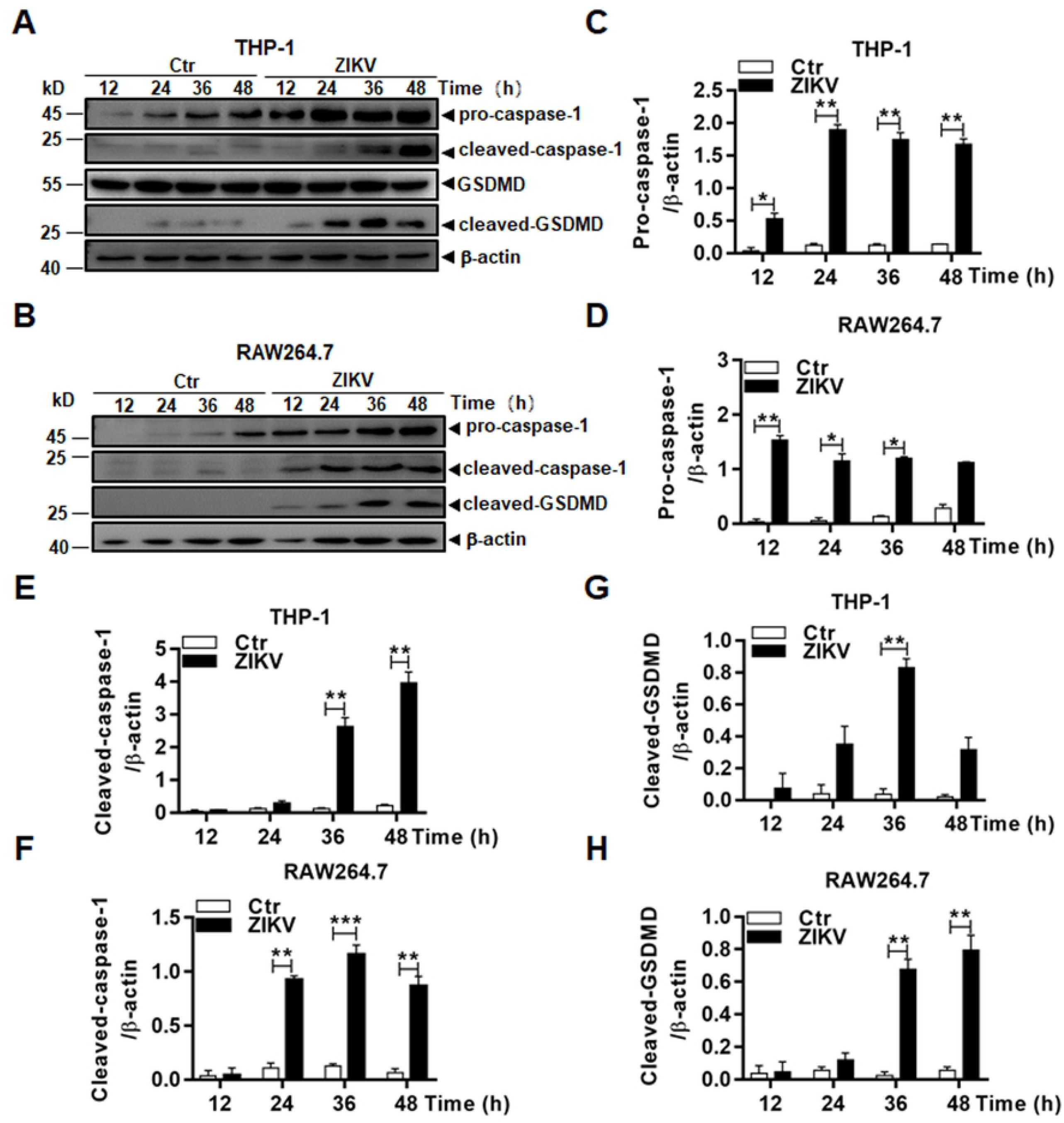
Inflammasome activation led to pyroptosis in ZIKV-infected monocytes. Cell lysates, prepared at various time points p.i. from THP-1 (**A**) and RAW264.7 (**B**) cells infected with 1 MOI ZIKV, were analysed by western blot analyses with antibodies for pro- or cleaved caspase-1 and GSDMD. Quantitative analyses of the grayscale values in the blots of the pro- and cleaved caspase-1 and GSDMD in infected and control THP-1 (**C, D**, & **F**) and RAW264.7 (**E, G**, & **H**) cells were shown. The experiments were performed in triplicates and the data were shown as mean+SD, analysed by unpaired Students t-test. *, P<0.05; **, P<0.01; ***, P<0.001.

### Upregulation of the Components for Inflammasome Formation and Its Activation in ZIKV-infected Monocytes

We examined the transcription and post-transcriptional processing of the inflammasome components in infected cells. RNA transcripts of the pro-caspase-1 (Figure 5A & D), NLRP3 (Figure 5B & C), and ASC (Figure 5 C & F) genes increased significantly over the time p.i. in both human and murine monocytes infected with ZIKV. We also examined the transcripts of pro-IL-1β and pro-IL-18 and found their RNA copy numbers increased as well in infected THP1 (Figure 6 A-B) and RAW264.7 cells (Figure 6 C-D). We confirmed that IL-1β and IL-18 were released to the culture medium of the infected THP-1 (Figure E-F) and RAW264.7 (Figure G-H) cells by ELISA.

**Figure 5.**
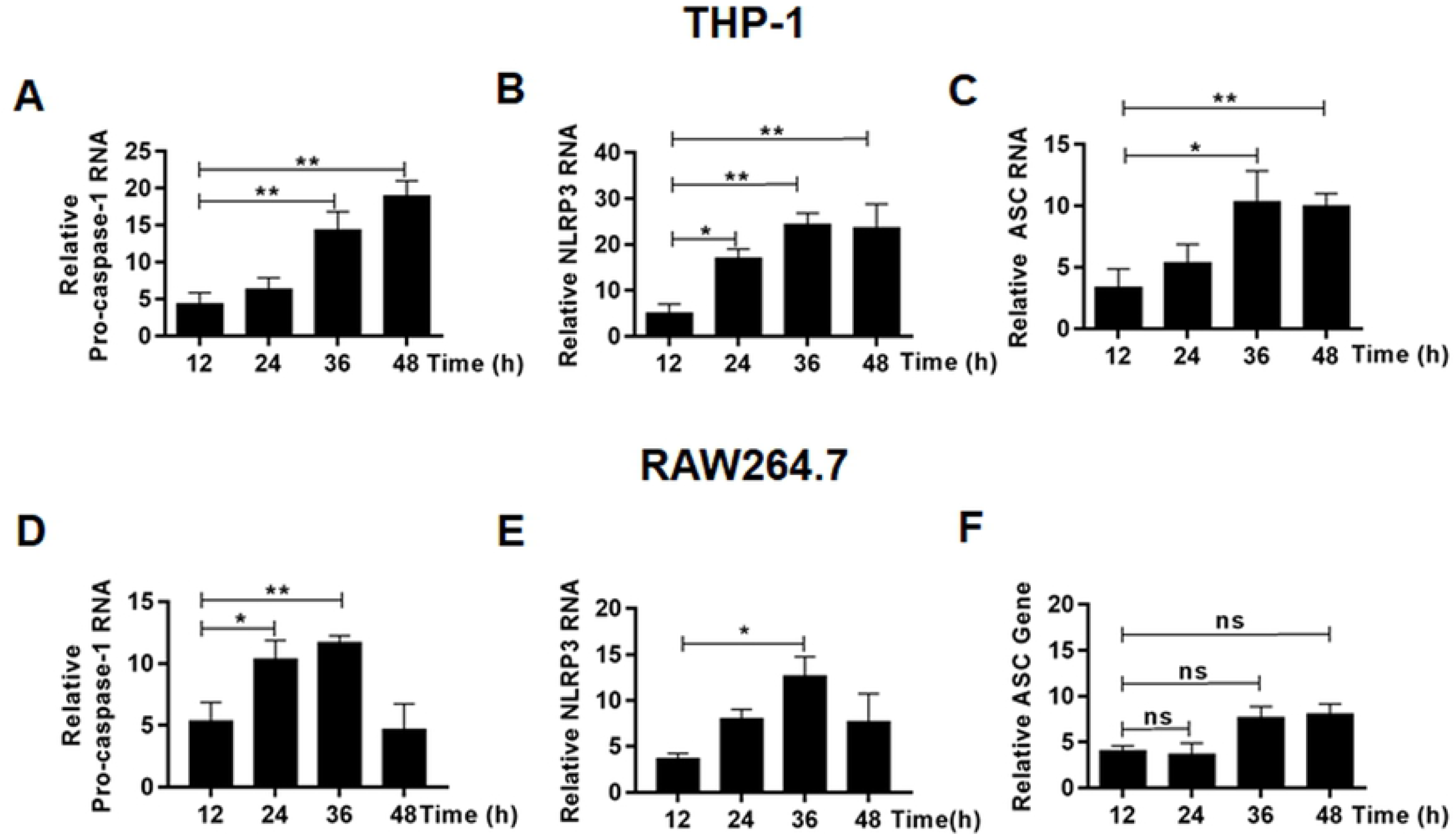
Upregulation of the inflammasome components in ZIKV-infected monocytes. Total RNA was prepared at various time points p.i. from THP-1 (**A-C**) and RAW264.7 (**D-F**) cells infected with 1 MOI of ZIKV for realtime RT-PCR to measure mRNA transcript numbers of pro-caspase-1 (**A** & **D**), NLRP3 (**B** & **E**), ASC (**C** & **F**). The experiments were performed in triplicates and the data were shown as mean+SD, analysed by unpaired Students t-test. *, P<0.05; **, P<0.01; ***, P<0.001. ns, no significance.

**Figure 6.**
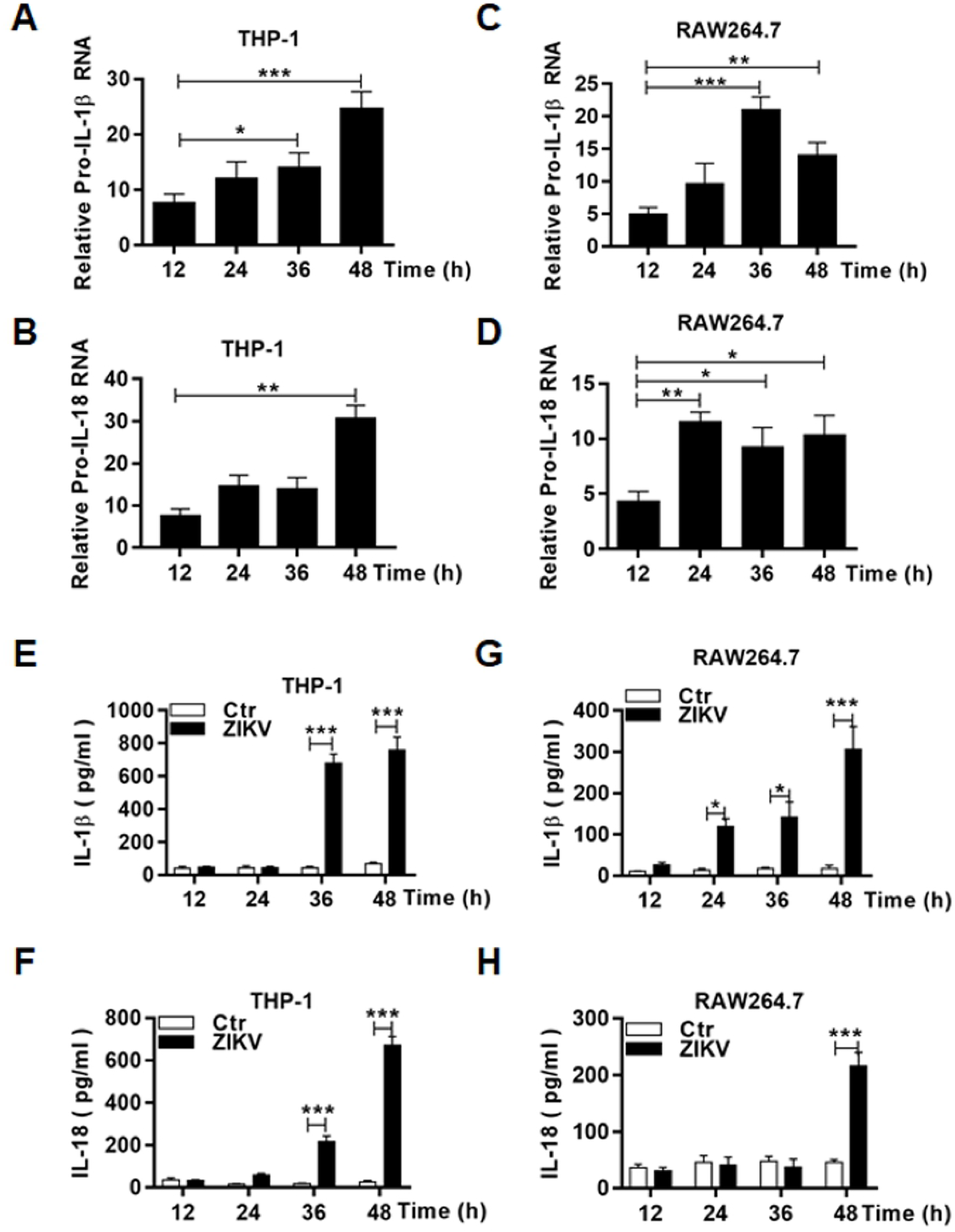
Transcriptional upregulation and secretion of IL-1β and IL-18 in ZIKV-infected human and murine monocytes. Total RNA was prepared at various time points p.i. from THP-1 (**A-B**) and RAW264.7 (**C-D**) cells infected with 1 MOI of ZIKV for realtime RT-PCR to measure mRNA transcript numbers of pro-IL-1β (**A** & **C**) and IL-18 (**B**-**D**) genes. Culture medium was sampled at various time points p.i. from ZIKV-infected THP-1 (**E-F**) and RAW264.7 (**G-H**) cells for measurement of secreted IL-1β (**E** & **G**) and IL-18 (**F** & **H**) by ELISA. The experiments were performed in triplicates and the data were shown as mean+SD and analysed by unpaired Students t-test. *, P<0.05; **, P<0.01; ***, P<0.001. ns, no significance.

### Inflammasome Activation Led to Cell Death Induced by ZIKV in monocytes

Pyroptosis can be triggered by processing of pro-caspase 1 via a canonical inflammasome activation, or processing of pro-caspase 4, 5, or 11 via a non-canonical approach. To confirm the mechanism for pyroptosis that occurred in monocytes infected with ZIKV, we chose to use Belnacasan, or VX-765, a specific caspase-1 inhibitor to pre-treat the cells prior to viral infection. Cell lysates were prepared at various time points p.i. for western blot analyses. As shown in Figure 7, cleaved caspase-1 did not appear or the level of cleaved caspase-1 was greatly reduced in VX765-treated monocytes, in comparison to the cells without treatment, indicating that VX765 effectively suppressed the activation of pro-caspase-1 in infected THP-1 (Figure 7A) and RAW264.7 (Figure 7B) cells. We examined the morphology of infected THP-1 and RAW247.1 cells at 48 hrs p.i. with or without the treatment of VX765. The amount of the cells with sick morphology increased significantly in both THP-1 (Figure 7C) and RAW247.1 (Figure 7D) cells, which were pre-treated with VX756, indicating that pro-caspase-1 cleavage or activation of inflammasomes was critical to triggering pyroptosis in ZIKV-infected monocytes.

**Figure 7.**
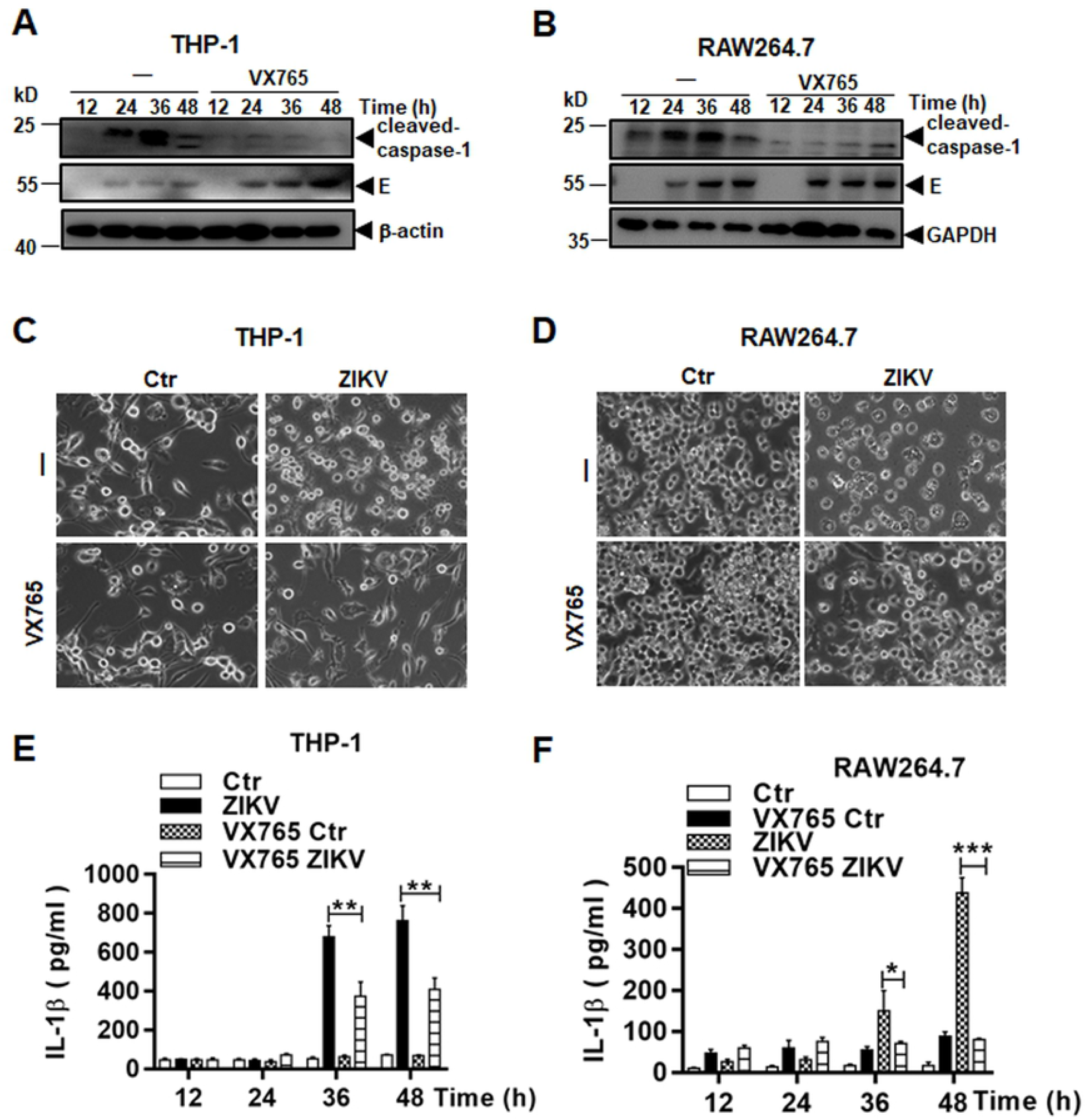
Pyroptosis was dependent on caspase-1 in ZIKV-infected human and murine monocytes. Cell lysates were prepared at various time points p.i. from ZIKV-infected THP-1 (**A**) and RAW264.7 (**B**) cells which were untreated or pre-treated with VX765, a caspase-1 inhibitor, at 20μM. The lysates were analysed by western blot analyses with an anti-cleaved caspase-1 antibody. Morphological changes were observed on ZIKV-infected THP-1 (**C**) and RAW264.7 (**D**) cells, which were untreated or pre-treated with VX765, under a light microscope (Magnification x40). Culture medium was sampled at various time points p.i. from ZIKV-infected THP-1 (**E**) and RAW264.7 (**F**) cells, which were untreated or pre-treated with VX765, for measurement of secreted IL-1β by ELISA. The experiments were performed in triplicates and the data were shown as mean+SD and analysed by unpaired Students t-test. *, P<0.05; **, P<0.01; ***, P<0.001.

To ascertain the inflammasome activation to be inhibited, we detected the secretion of IL-1β in infected monocytes, pre-treated with or without VX765. The increase of IL-1β in infected cells was significantly suppressed at 36 and 48 hrs p.i. in VX765 pre-treated cells, compared to the cells without the treatment (Figure 7E-F).

We further used a small hairpin RNA (shRNA) approach to confirm the role of the inflammasome activation involved in the cell death. Three shRNA molecules, targeting pro-caspase-1, were tested in THP-1 cells to knock down pro-caspase-1 transcription. The expression of pro-caspase-1 was suppressed in the cell lines, which were selected and expanded, as shown in Figure 8A. The caspase-1 knockdown (KD) cell lines were infected with ZIKV and cell lysates were prepared at various time points p.i. for western blot analysis, which showed little expression of pro-caspase-1, barely detectable cleaved caspase-1, and no detection of cleaved GSDMD, a pattern distinct from those in the control cells infected with ZIKV (Figure 8B). Cell viability was measured by an MTT assay and significant cell death occurred in ZIKV-infected cells at 36 and 48 hrs p.i. However in pro-caspase-1 KD cells, the cell death was reversed significantly in the absence of pro-caspase-1 (Figure 8C), indicating that the pyroptosis, that occurred in ZIKV-infected monocytes, was caspase-1 dependent via a canonical pathway.

**Figure 8.**
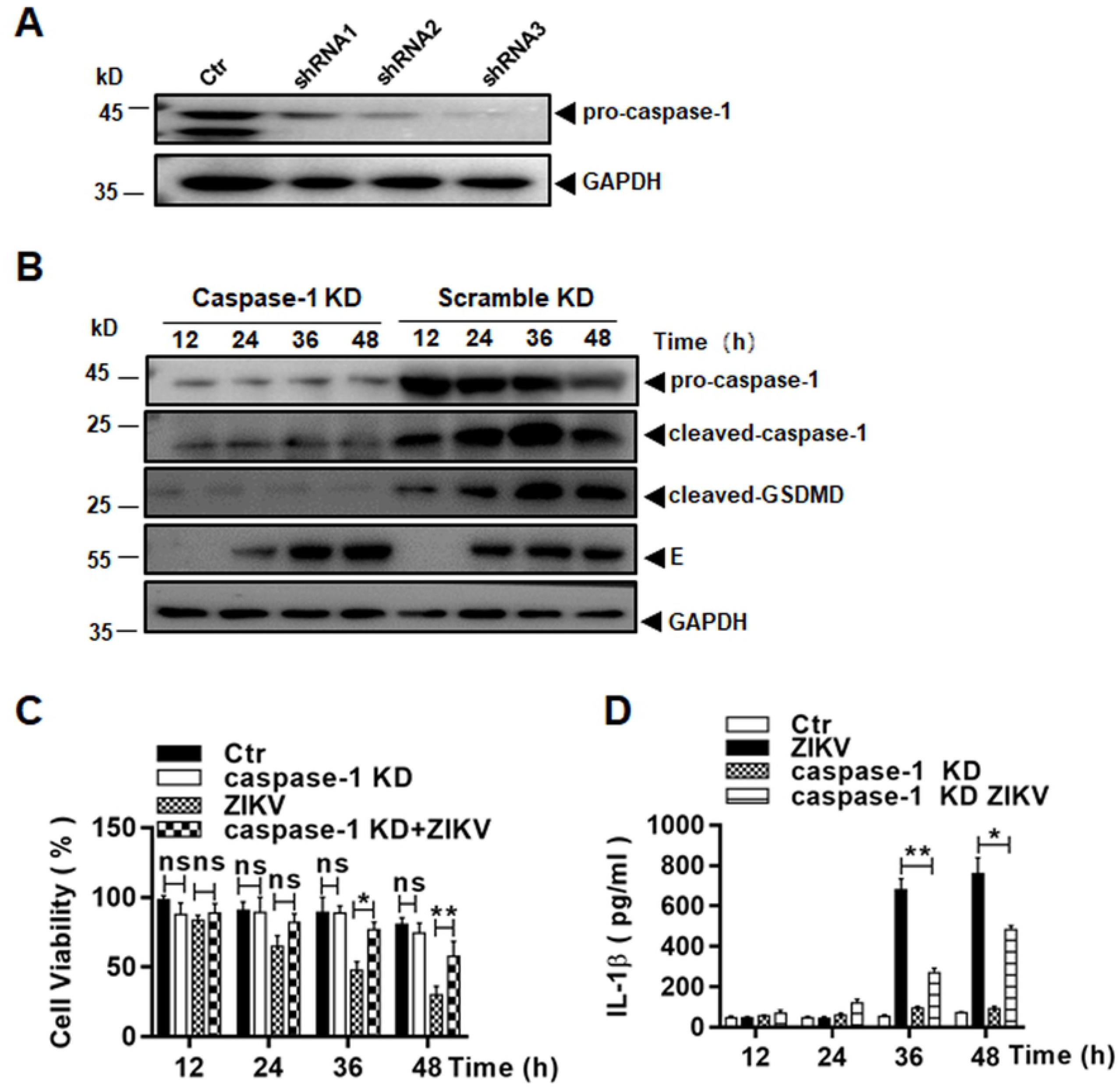
Pyroptosis was suppressed in ZIKV-infected monocytes with pro-caspase-1 knockdown. (**A**) Knockdown of pro-caspase-1 in THP-1 cells. Cell lysates were prepared from lentiviral vector-transduced THP-1 cell lines, expressing shRNA1, 2, or 3 targeting mRNA of pro-caspase-1, for western blot analyses with pro-caspase-1 antibody to examine the knockout (KD) efficacy. (**B**) Inhibition of pro-caspase-1 and GSDMD processing in pro-caspase-1 KD cells. Both pro-caspase-1 or scramble shRNA KD cells were infected with ZIKV and cell lysates, prepared at various time points p.i., were subjected to western blot analyses with antibodies for pro-caspase-1, cleaved caspase-1 and GSDMD. (**C**) Cell viability was assessed in pro-caspase-1 or scramble shRNA KD THP-1 cells infected with or without ZIKV at various MOIs by an MTT assay. (**D**) Culture medium was sampled at various time points p.i. from pro-caspase-1 or scramble shRNA KD cells infected with or without ZIKV for measurement of secreted IL-1β by ELISA. The experiments were performed in triplicates and the data were shown as mean+SD and analysed by unpaired Students t-test. *P, <0.05; **, P<0.01. ns, no significance.

To ascertain the effect on the inflammasome activation by pro-caspase-1 KD in the cells, we measured the increase of IL-1β which was significantly suppressed at 36 and 48 hrs p.i., compared to the infected control cells without pro-caspase-1 KD, confirming that the pro-caspase-1 KO was sufficient in suppressing the inflammasome activation in ZIKV-infected monocytes (Figure 8D).

### The NLRP3 Inflammasome Was Essential to ZIKV-induced Pyroptosis

As described earlier, NLRP3 was transcriptionally upregulated in human monocytes infected with ZIKV. To confirm the role of the NLRP3 inflammasome activation in pyroptosis, we carried out the shRNA knockdown of NLRP3 in THP-1 cells. Three shRNA molecules, targeting NLRP3, were tested in THP-1 cells to knock down NLRP3 transcription. The cell lines were selected and examined for their knockdown efficacy. The result with efficient knockdown of NLRP3 was shown in Figure 9A. The NLRP3 knockdown (KD) cells were infected with ZIKV and cell lysates were prepared for western blot analysis. In infected control cells, pro-caspase-1 was processed and cleaved GSDMD was produced, but in infected KD cells, pro-caspase-1 was barely processed and cleaved GSDMD was not produced (Figure 8B), indicating that the pyroptosis, triggered in infected monocytes, was dependent on the NLRP3 inflammasome activation, a canonical approach, leading to processing of pro-caspase 1.

**Figure 9.**
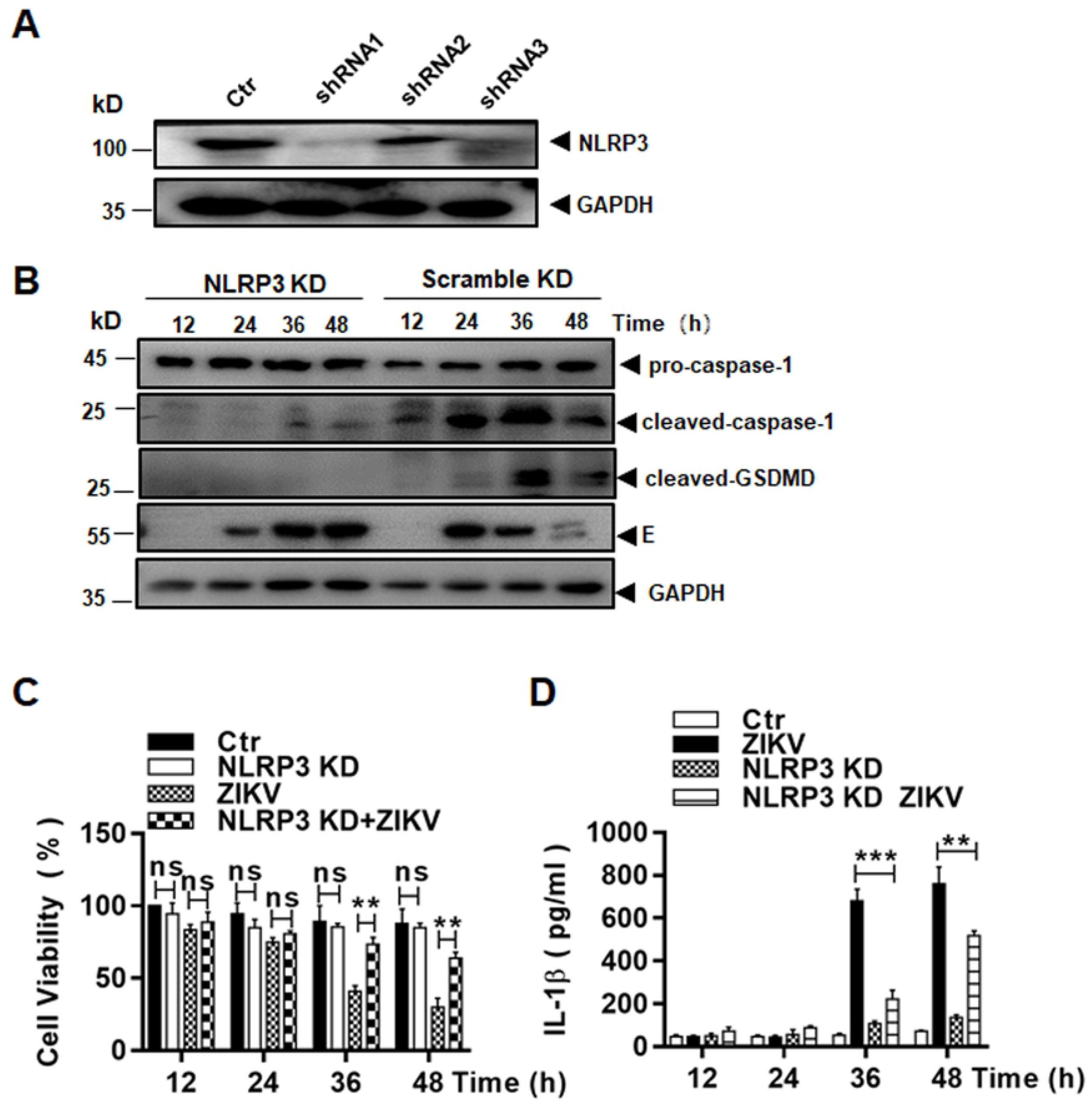
Pyroptosis triggered in ZIKV-infected monocytes were dependent on the NLRP3 inflammasome activation. (**A**) Knockdown of NLRP3 in THP-1 cells. Cell lysates were prepared from lentiviral vector-transduced THP-1 cell lines, expressing shRNA1, 2, or 3 targeting mRNA of NLRP3, for western blot analyses with an NLRP3 antibody to examine the knockdown (KD) efficacy. (**B**) Inhibition of pro-caspase-1 and GSDMD processing in NLRP3 KD cells. Both NLRP3 or scramble shRNA KD cells were infected with ZIKV and cell lysates, prepared at various time points p.i., were subjected to western blot analyses with antibodies for pro-caspase-1, cleaved caspase-1 and GSDMD. (**C**) Cell viability was assessed in NLRP3 or scramble shRNA KD THP-1 infected with or without ZIKV at various MOIs by an MTT assay. (**D**) Culture medium was sampled at various time points p.i. from NLRP3 or scramble shRNA KD cells, infected with or without ZIKV, for measurement of secreted IL-1β by ELISA. The experiments were performed in triplicates and the data were shown as mean+SD and analysed by unpaired Students t-test. **, P<0.01; ***, P<0.001. ns, no significance.

Cell viability was measured in NLRP3 KD cells after infection with MTT assay. As shown in Figure 9C, significant cell death occurred at 36 and 48 hrs p.i. in infected control cells but the cell death was effectively suppressed in the NLRP3 KD cells infected with ZIKV.

To confirm that the NLRP3 inflammasome activation was impaired in the NLRP3 KD cells, we further measured the secretion of IL-1β in infected control and NLRP3 KD cells by ELISA, which showed that the increase of IL-1β secretion in the culture medium was significantly reduced in the NLRP3 KD cells (Figure 9D). In sum, these data demonstrate that the activation of the NLRP3 inflammasome led to pyroptosis, which was dependent on activated caspase-1 in ZIKV-infected monocytes

## DISCUSSION

ZIKV causes asymptomatic or mild infections, which are self-limited in most adults, indicating that host immunity can contain the infection effectively in most adults. However, ZIKV can dissimilate through bloodstream to the placenta in some pregnant women and penetrate the blood-placenta barrier (BPB) into fetus, in which the virus invades the neural tissues due to its neurotropism. What role monocytes play in facilitating the virus to penetrate the BPB barrier and eventually infect the fetal brain remains to be investigated. In this study, we confirmed that both human and murine monocytes were susceptible to ZIKV, and a productive replication led to cell death. The cell death could be observed at 12 hrs p.i., and continued to eventually deteriorate the whole cell culture, suggesting that ZIKV caused a lytic infection in monocytes in addition to released proinflammastory cytokines and chemokines as shown in this and previous studies. We were able to identify that apoptotic process was triggered upon ZIKV infection, which also occurs in monocytes infected with many types of viruses (25-29).

Previous studies have shown that ZIKV infection activates the NLRP3 inflammasome, which leads to processing of pro-caspase-1 and secretion of IL-1β in infected monocytic cell lines, PBMC, or monocyte-derived macrophages (13-15, 17). Interestingly, none of these studies have pursued to observe or detect pyroptosis induced in infected monocytes or macrophages. In this report we showed the occurrence of pyroptosis, in addition to apoptosis, in ZIKV-infected monocytes. As reported previously, we showed that pro-caspase-1 was cleaved and the secretion of cytokines IL-1β and IL-18 increased, indicating that inflammasomes were activated in monocytes. Cleaved GSDMD, an executor of pyroptosis, was further detected, suggesting that a considerate amount of cell death in ZIKV-infected monocytes was attributed to pyroptosis, which could be suppressed in the cells, pre-treated with an inhibitor for caspase-1 or shRNA to knockdown caspase-1. We finally showed that the NLRP3 inflammasome activation was required for pyroptosis via a canonical approach in ZIKV-infected monocytes, which was dependent on caspase-1 in infected human and murine monocytes.

Monocytes and macrophages are important in viral infections. As haemopoietic cells originated in bone marrow, monocytes comprise about 10% of blood leukocytes. They are released into peripheral circulation and live for a few days in blood vessels of the body. Monocytes penetrate through the wall of vessels and reside in the tissues and organs, through chemotaxis when microbial infection occurs, and differentiate into macrophages. Monocytes are susceptible to many viruses in various families. Human monocyte-derived macrophages can be infected by Coxsackieviruses CV-B4(30), which, however, can poorly infect human monocytes, unless a non-neutralizing anti-CV-B4 IgG is present (31). Replication of neurovirulent poliovirus strains in monocytes was associated to their pathogenesis in the central nervous system (32). Monocytes and macrophages play critical roles in HIV transmission, viral spread early in the host, and being a reservoir of virus throughout infection (8).

Circulating monocytes are the primary cellular target of ZIKV infection in humans. Not only can monocytes be infected dominantly in PBMC by ZIKV in vitro, but the virus can also be found in monocytes collected during acute illness from Zika patients (11, 33). ZIKV viral RNA can be detected in the monocytes longer than in the serum of infected patients, indicating that monocytes could serve as a virus reservoir during the infection (33). Interestingly, infection of monocytes by ZIKV leads to cell expansion to become more intermediate or non-classical type by expressing CD16 (11, 33). Moreover, monocytes infected with ZIKV tend to secrete IL-10, an immunosuppressive cytokine, which could skew the host immune response during the early stage of pregnancy (11). These reports collectively support that monocytes may play a complicated role in viral pathogenesis on ZIKV spread and neuropathogenesis. In this sense, programed cell death in ZIKV-infected monocytes, not reported previously to our knowledge, may be beneficiary to the host as a protective defence by eliminating the virus reservoir in the host.

Programed cell death can be triggered in monocytes infected with other flaviviruses. It has been observed that dengue virus (DENV) can induce apoptosis in infected monocytes, which is related to increased TNF-α induction (29). In fact all four serotypes of DENV can induce apoptosis in human monocytes, dependent on activation of caspase 7, 8, and 9 (34). On the other hand, DENY can also trigger pyroptosis, which is associated with an activation of caspase-1 and release of IL-1β (35). In this report our data demonstrate that a concomitant apoptosis and pyroptosis were induced in ZIKV-infected human and murine monocytes. It was reported that the NLRP3 inflammasome activation, triggered by ZIKV infection in monocytes, promotes the cleavage of cGAS, resulting in the inhibition of initiating type I IFN signaling, and enhances viral replication (13). We believe that subsequent pyroptotic cell death, caused by the NLRP3 inflammasome activation, or caspase-3 dependent apoptosis as shown in our data may help shorten or clear viral replicative locales and rid the viral carrier in the host. The programed cell death may be also significant in preventing the virus from spreading to the placenta for infecting fetal chorionic villi during pregnancy.

Little is known about why an infected cell chooses one way or another to die. In contrast to apoptosis, pyroptosis and necroptosis are inflammatory, leading to massive damage of involved tissues. Cross-talk occurs between signaling pathways of the programmed cell deaths, which may provide a mechanism for regulating cell fate. We have identified that monocytes are programmed to pyroptosis and apoptosis in response to ZIKV infection. Since ZIKV nonstructural protein NS5 is required for NLRP3 activation (15), the pyroptosis is therefore very likely to be triggered by viral protein NS5 in monocytes. We cannot determine the exact mechanism about how apoptosis is induced but very likely the induction of TNF-α (36) in infected monocytes plays an important factor in activating the cascade of caspases leading to the processing of pro-caspase-3. We have no clue at this stage whether these programed cell death pathways may regulate each other in ZIKV-infected monocytes. Even though caspase-3 can cleave GSDMD, leading to pyroptosis (37), we showed in this study that ZIKV-triggered pyroptosis in monocytes was caspase-1 dependent via a canonical approach. Our data in this report may help us further understand the complicated functions how monocytes are involved in viral pathogenesis in ZIKV-infected humans.

## ACKNOWLEDGEMENTS

We are grateful to Dr. Shibo Jiang, Fudan University, for the Zika virus strain used in the study.

## CONFLICT OF INTEREST

The authors declare that they have no conflicts of interest with the contents of this article.

## AUTHOR CONTRIBUTIONS

CXW and ZX conceived and coordinated the study. CXW, YFY, and CFG designed, performed and analyzed the experiments shown in Figures 1 through 9. XQ provided reagents, technical assistance and contributed to completion of the studies. CXW, CJC and ZX wrote the paper. All authors reviewed the results and approved the final version of the manuscript.

## DATA AVAILABILITY

The datasets used and/or analyzed during the current study are available from the corresponding author on reasonable request.

## SUPPLEMENTAL INFORAMTION

**Figure S1.** Activation of Caspases and Phosphorylation of RIPKs in ZIKV-infected Monocytes. THP-1 cells were infected with ZIKV and cell lysates were prepared at various time points p.i. for western blot analyses with antibodies for pro- and cleaved caspase-3, PARP (**A**) and RIPK1, RIPK3, and MLKL (**B**).

## Notes

### Competing Interest Statement

The authors have declared no competing interest.

